# An improved TurboID pipeline in *C. elegans* by biochemical depletion of endogenously biotinylated carboxylases

**DOI:** 10.1101/2022.05.22.492627

**Authors:** Murat Artan, Markus Hartl, Weiqiang Chen, Mario de Bono

## Abstract

Proximity-dependent protein labeling provides a powerful *in vivo* strategy to characterize the interactomes of specific proteins. We previously optimized a proximity labeling protocol for *C. elegans* using the highly active biotin ligase TurboID. A significant constraint on the sensitivity of TurboID is the presence of abundant, endogenously biotinylated proteins that take up bandwidth in the mass spectrometer, notably carboxylases that use biotin as a co-factor. In *C. elegans*, these comprise POD-2/acetyl-CoA carboxylase alpha, PCCA-1/propionyl-CoA carboxylase alpha, PYC-1/pyruvate carboxylase and MCCC-1/methylcrotonyl-CoA carboxylase alpha. We developed ways to remove these carboxylases prior to streptavidin purification and mass spectrometry, by engineering their corresponding genes to add a C-terminal His_10_ tag. This allows us to deplete them from *C. elegans* lysates using immobilized metal affinity chromatography (IMAC). To demonstrate the method’s efficacy, we use it to expand the interactome map of the presynaptic active zone protein ELKS-1. We identify many known active zone proteins, as well as previously uncharacterized potentially synaptic proteins. Our approach provides a quick and inexpensive solution to a common contaminant problem in biotin- dependent proximity labeling. The approach may be applicable to other model organisms and will enable deeper and more complete analysis of interactors for proteins of interest.

## Introduction

Protein-protein interactions underlie most biological processes. Methods that highlight such interactions can provide key insights into basic molecular mechanisms and help target strategies for therapeutic interventions (1). One such method is proximity-dependent protein labeling, which can both characterize protein-protein interaction networks and define the proteomes of subcellular structures (1, 2). The proximity labeling enzymes most widely used in living cells and organisms are currently engineered variants of the *E. coli* biotin ligase BirA, such as TurboID, the ascorbate peroxidase APEX and its derivatives, and horseradish peroxidase HRP (1–3). These enzymes all release activated biotin, which reacts with amine groups of exposed lysine residues in the vicinity (4–6).

In living organisms biotin is a critical cofactor of carboxylation reactions. Biotin-dependent carboxylases, notably acetyl-CoA carboxylase (ACC), propionyl- CoA carboxylase (PCC), 3-methylcrotonyl-CoA carboxylase (MCC), and pyruvate carboxylase (PC) are widespread in nature, and important for the metabolism of fatty acids, carbohydrates and amino acids. In mammals, which do not synthesize or store biotin, biotin deficiency causes severe syndromes, including ataxia and neurological dysfunction (7, 8). Biotin is covalently linked to the carboxylase apoproteins by holocarboxylase synthetase (HLCS) (7). The nematode *C. elegans* has one ortholog of HLCS, BPL-1 (biotin protein ligase 1), which biotinylates the carboxylases MCCC-1/MCCC (methylcronotoyl coenzyme A carboxylase), PCCA- 1/PCCA (propionyl coenzyme A carboxylase alpha subunit), POD-2/ACACA (acetyl coenzyme A carboxylase) and PYC-1/PC (pyruvate carboxylase) (9). These four endogenously biotinylated carboxylases, from here on referred to collectively as MP3, and their orthologs, dominate the biotinylated proteome of many organisms, including worms, flies, mice, and humans. MP3 proteins provide by far the most abundantly detected peptides when streptavidin affinity-purified samples are analyzed by mass spectrometry (10–12).

High abundance proteins (HAPs) in biological samples limit detection of low abundance proteins (LAPs) by mass spectrometry (MS) (13). Significant effort has been spent developing ways to deplete HAPs from different samples, to increase detection of LAPs by MS. For example, collagenase depletion of collagen significantly increases the number and variety of proteins detected from extracellular matrix (ECM)-derived protein extracts (14). Albumins, immunoglobulins, fibrinogen, transferrin and haptoglobulin comprise more than 90% of the plasma proteome and therefore, mask the detection of LAPs. Depleting albumins and immunoglobulins by affinity chromatography prior to MS analysis makes it possible to identify many LAPs in blood plasma (15). EDTA-functionalized nanoparticles effectively double the number of proteins detected in urine samples (16). Extraction protocols that favor recovery of LAPs over HAPs (beta-conglycinin and glycinin) from soybean seeds allow more LAPs to be identified (17).

Proximity labeling has been applied extensively in cultured cells but its use in *C. elegans* is in its infancy. Overexpressing a patronin::TurboID fusion protein in the worm gut successfully highlighted the interactome of this microtubule-binding protein (18). Fusing TurboID to an anti-GFP nanobody allowed tissue-specific analysis of the interactomes of the GFP-tagged centrosomal proteins PLK-1 and SPD-5 (19). We identified interactors of the presynaptic active zone protein ELKS-1 by knocking in a TurboID::mNeongreen cassette at the *elks-1* locus (10). We also found that knock- ins that express TurboID fusions at endogenous levels enhance the number of interacting partners identified compared to transgenically overexpressing the corresponding fusion. However, for proteins of interest (POI) expressed at low levels, or in a small number of cells, detecting biotinylated proximal interactors by MS becomes difficult due to the abundant co-purifying MP3 proteins. Here, we devise a way to deplete MP3 proteins prior to streptavidin-based affinity purification, thus increasing the signal/noise ratio for TurboID-based proximity labeling in *C. elegans*. Using ELKS-1 as our POI we show that depleting MP3 enables many more known or putatively synaptic proteins to be identified compared to analysis of undepleted controls.

## Results

To increase the signal/noise ratio in biotin-dependent proximity labeling we first sought to prevent endogenous MP3 biotinylation. To achieve this, we tried to deplete BPL-1, the enzyme that biotinylates MP3. BPL-1 is required for efficient *de novo* fatty acid biosynthesis and deleting *bpl-1* results in maternal-effect embryonic lethality (9). To conditionally disrupt *bpl-1*, we knocked in an auxin-inducible degron (AID) just upstream of the *bpl-1* termination codon. Incubating *bpl-1::AID* worms with auxin from the egg, L1 or L2 stages arrested their growth. By contrast, worms transferred to auxin plates from the mid-L3 stage and onward grew to adulthood. However, when we harvested these animals as young adults and probed Western blots of their extracts with streptavidin, we did not observe reduced biotinylation of MP3 (Fig. S1A). Incubating *bpl-1::AID* worms on auxin plates for longer periods, for 48 or 64 hours post-YA stage, depleted biotinylated POD-2 but not the other three proteins (Fig. S1B).

Since knocking down BPL-1 only had marginal effects on MP3 biotinylation, we sought to deplete each carboxylase individually. Using Crispr/Cas9-mediated genome editing, we tagged each protein with an AID at its C terminus. *mccc-1* and *pcca-1* only express one isoform of 73.7 and 79.7 kDa, respectively, while *pod-2* expresses three isoforms of 91.4, 230.6 and 242.6 kDa and *pyc-1* two isoforms of 67.8 and 129.3 kDa. We chose to tag the C termini because they are shared by all isoforms. We grew the *mccc-1::3XFLAG-AID; pcca-1::3XFLAG-AID; pyc-1::3XFLAG- AID* expressing animals (MP2-deg worms) on auxin plates for several generations but only achieved mild depletion of these 3 mitochondrial matrix proteins (Fig. S1C). Growing animals that expressed *pod-2::3XFLAG-AID* as well as MP2::AID (henceforth referred to as MP3-AID) on auxin plates caused embryonic or L1 lethality. Transferring MP3::AID worms from no auxin control plates to auxin plates depletes POD-2::AID within 5 hours and caused strong phenotypes, but did not alter levels of the other carboxylases (Fig. S1C). Presumably, a 5-hour incubation on auxin is insufficient to deplete MCCC-1, PCCA-1 and PYC-1 because they are localized in the mitochondrial matrix. Ubiquitination does occur in different mitochondrial compartments, and mitochondrial proteins can be degraded in a ubiquitin-proteasome system (UPS)-dependent manner (20). We therefore explored if targeting TIR1 to mitochondria can promote efficient degradation of MP3-AID proteins. We found, however, that F1 animals expressing TIR-1::BFP in mitochondria arrested at L1 stage, and we abandoned this approach after several unsuccessful attempts to recover edited worms.

Since we were unable to deplete all four carboxylases and their various isoforms satisfactorily using the AID system, we sought a different way to remove biotinylated carboxylases from worm extracts. Immobilized metal affinity chromatography (IMAC), using for example Ni-NTA resins, allows efficient purification of proteins bearing a polyhistidine tag on either terminus. IMAC separation offers several advantages: the interaction between the polyhistidine tag and Ni^2+^ resin is strong and stable in harsh environments; Ni^2+^-NTA resins have high protein binding capacity and can be regenerated; and the resins are relatively cheap (21). We decided to take an IMAC-based approach to deplete MP3 from *C. elegans* lysates (Fig. 1A). We engineered each of the genes encoding MP3 carboxylases to add His_10_ tags to their C termini (MP3-His). We then introduced transgenes that express either mNeongreen alone (*rab-3p::mNG*) or a TurboID::mNeongreen fusion throughout the nervous system (*rab-3p::TurboID::mNG*) into the MP3-His_10_ genetic background to optimize an MP3 depletion protocol. Incubating lysates with Ni-NTA resin efficiently deplete MP3 from the samples with minimal loss of other biotinylated proteins (Figure 1B). Subjecting the samples to two rounds of Ni-NTA resin-based depletion was not necessary to capture most of the His_10_-tagged proteins (Figure 1B); a second round of depletion did not significantly impact the MS analysis (data not shown). As expected, the Ni-NTA resin did not capture biotinylated proteins lacking a His_10_ tag, confirming to the specificity of the resin (Figure S2A).

**Figure 1.**
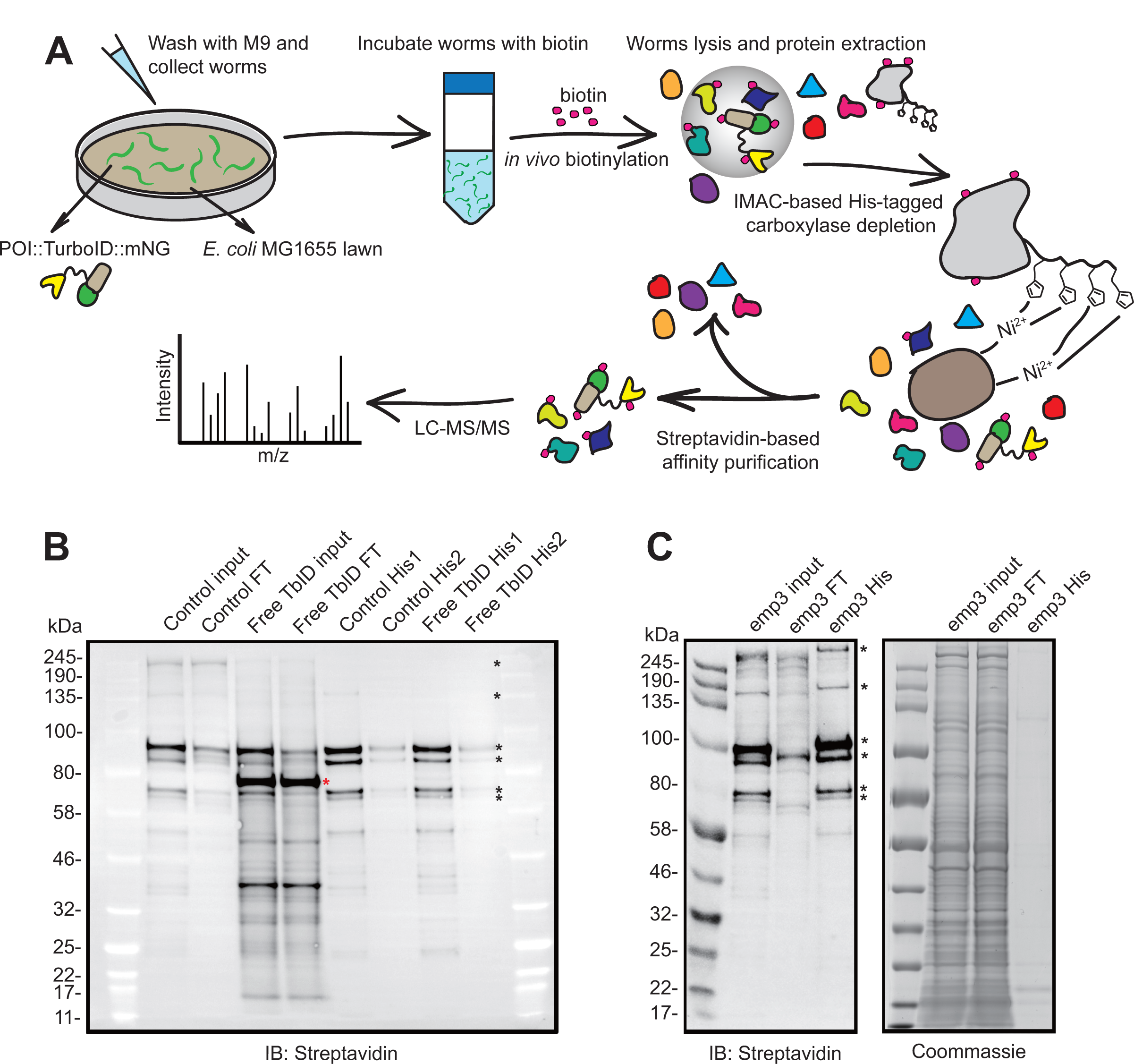
Optimizing TurboID-mediated proximity labeling in *C. elegans*. (**A**) Schematic overview of TurboID protocol, including depletion of His_10_-tagged endogenously-biotinylated carboxylases. (**B**) Western blot showing the efficiency of carboxylase depletion using a Ni-NTA resin and extracts from *C. elegans* expressing MCCC-1::His_10_; PCCA-1::His_10_; POD-2::His_10_; PYC-1::His_10_ (abbreviated as MP3-His). (**C**) Western blot analysis of extracts from animals expressing *elks- 1::TbID::mNG* in an MP3 background. One round of Ni-NTA depletion efficiently removes His_10_-tagged MP3. POI: protein of interest; input: whole lysate without carboxylase depletion; FT: flow through after carboxylase depletion; His1 and His2: Eluted His-tagged carboxylases captured by Ni-NTA resin after 1 (His1) or 2 (His2) rounds of His depletion. Control: *rab-3p::mNG; mp3-His*; Free TbID: *rab- 3p::TbID::mNG; mp3-His*; emp3: *elks-1::TbID::mNG; mp3-His.* Red asterisk: TurboID-mNeongreen; Black asterisks, endogenously biotinylated carboxylases.

We previously engineered the endogenous *elks-1* gene, which encodes a synaptic active zone protein, to tag ELKS-1 C-terminally with TurboID–mNeongreen. Affinity purifying extracts made from these animals with streptavidin enabled us to identify *bona fide* synaptic proteins using MS (10). To benchmark our MP3 depletion strategy, we probed the ELKS-1 interactome further. Western blot data showed efficient depletion of His_10_-tagged MP3 from the samples that we submitted for MS analysis (Fig. 1C). HPLC-UV analysis of peptides obtained from on-bead digestion following streptavidin purification revealed an approximately tenfold decrease in total peptide concentration in samples depleted of MP3-His_10_ compared to non-depleted controls (Fig. S2C). To match the amount of proteins analyzed across samples, we injected 5% of the undepleted samples and 50% of the depleted samples into the reverse phase HPLC column upstream of the MS. We also loaded 5% of the depleted samples to the column for comparison. LC-MS/MS analysis cumulatively identified 2430 proteins present in wild-type, depleted or undepleted ELKS-1 samples (Table S1).

Using the amica proteomics data analysis platform (22), we found that 180 proteins were enriched and 36 reduced (log_2_ ≥1.5 fold change threshold, adjusted p≤0.05) in undepleted ELKS-1::TbID samples compared to wild-type (Fig. 2A). MP3-depleted ELKS-1::TbID 50% and 5% loaded samples were further enriched for 214 and 133 proteins and reduced for 19 and 81 proteins, respectively compared to undepleted ELKS-1::TbID samples, (Fig. 2B, C, D). Although many proteins annotated on Wormbase (www.wormbase.org) to be at synapses were enriched in both depleted and undepleted lists, enrichment was significantly higher in the depleted list (Fig. 2E, S3A, S3B). Most, although not all, synaptic proteins already enriched in the undepleted list were further enriched in the depleted list, although, to a lesser extent than proteins that could not be identified without depletion (Fig. 2E).

**Figure 2.**
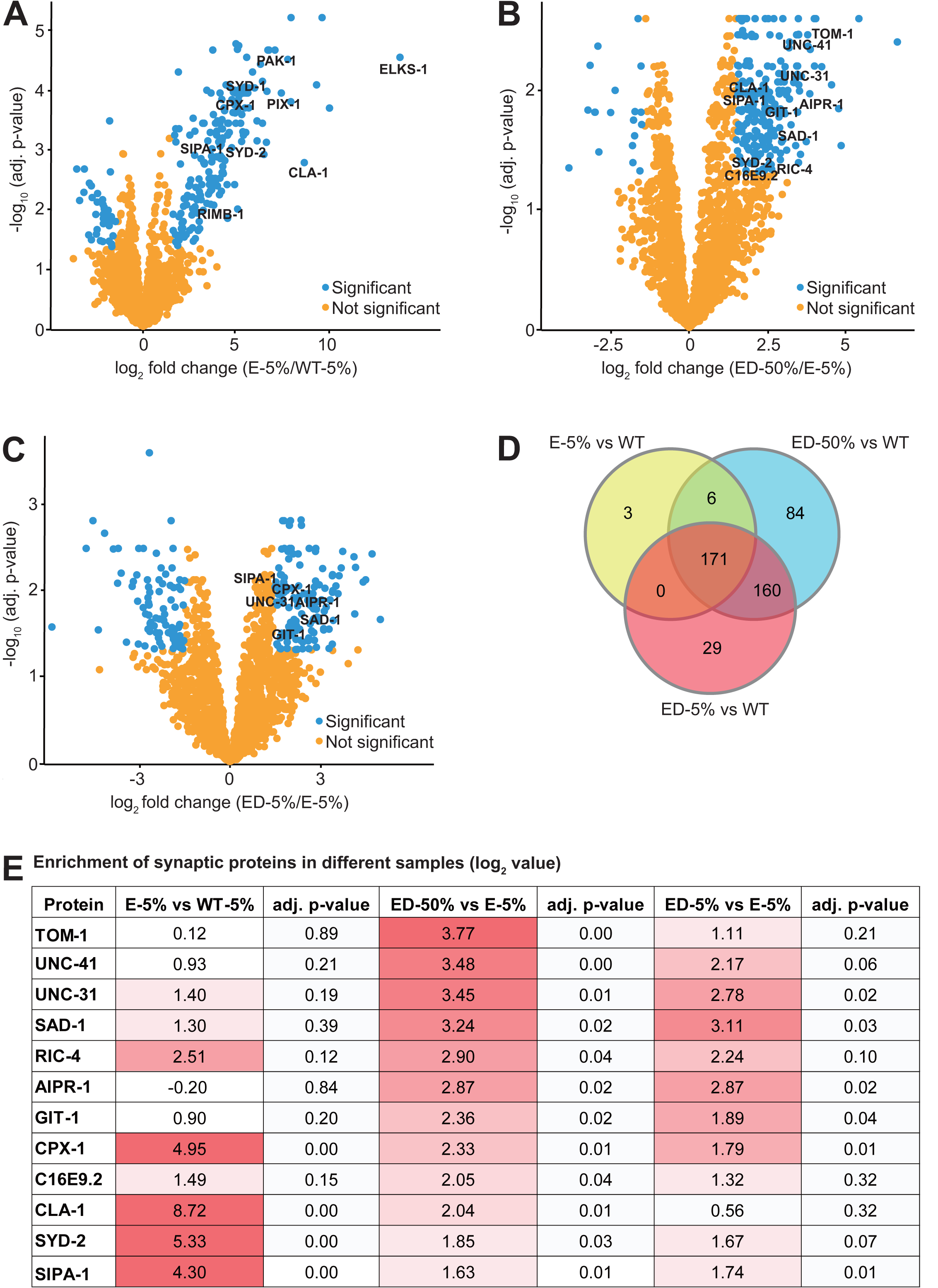
Interactome analysis of the pre-synaptic active zone protein ELKS-1 by comparing carboxylase depleted versus undepleted samples. (**A**) Significantly enriched proteins in ELKS-1 undepleted samples compared to wild-type (log_2_ ≥1.5 fold change threshold, adj. p value ≤0.05). As expected, many synaptic proteins are highly enriched. (**B** and **C**) Significantly enriched proteins when data obtained from analysis of 50% (**B**) or 5% (**C**) of depleted samples are compared to 5% of an undepleted ELKS-1 samples (log_2_ ≥1.5 fold change threshold, adj. p value ≤0.05). (**D**) Venn diagram highlighting number of significantly enriched proteins between samples. See also Fig. S3A. (**E**) Table showing log_2_ fold enrichment and adjusted p- value of synaptically annotated proteins. WT: *mp3-His_10_*; E: *elks-1::TbID::mNG* carboxylase undepleted; ED: *elks-1::TbID::mNG; mp3-His_10_* carboxylase depleted.

As well as enrichment of many known synaptic proteins, we identified uncharacterized proteins enriched in the 5% and 50% ELKS-1::TbID depleted samples that were not identified in the undepleted samples (Table S1, Fig. 3A, B and C). Among these proteins, F59C12.3 stood out due to the low number of spectral counts in the 3 replicates of undepleted ELKS-1–TbID samples (Table S1, Fig. 3A, B and C). We expressed mNeongreen translational fusion transgenes for this protein under the control of the DA9 neuronal promoter *itr-1*, and showed that it co-localized with ELKS-1::mScarlet at pre-synaptic active zones where DA9 synapses with neuromuscular junctions (23) (Fig. 3D). Altogether, our findings suggest that depleting MP3 in *C. elegans* lysates significantly improves the sensitivity of proximity-labeling using TurboID. The biochemical depletion approach we take may be applicable to other model organisms to enhance characterization of protein interaction networks.

**Figure 3.**
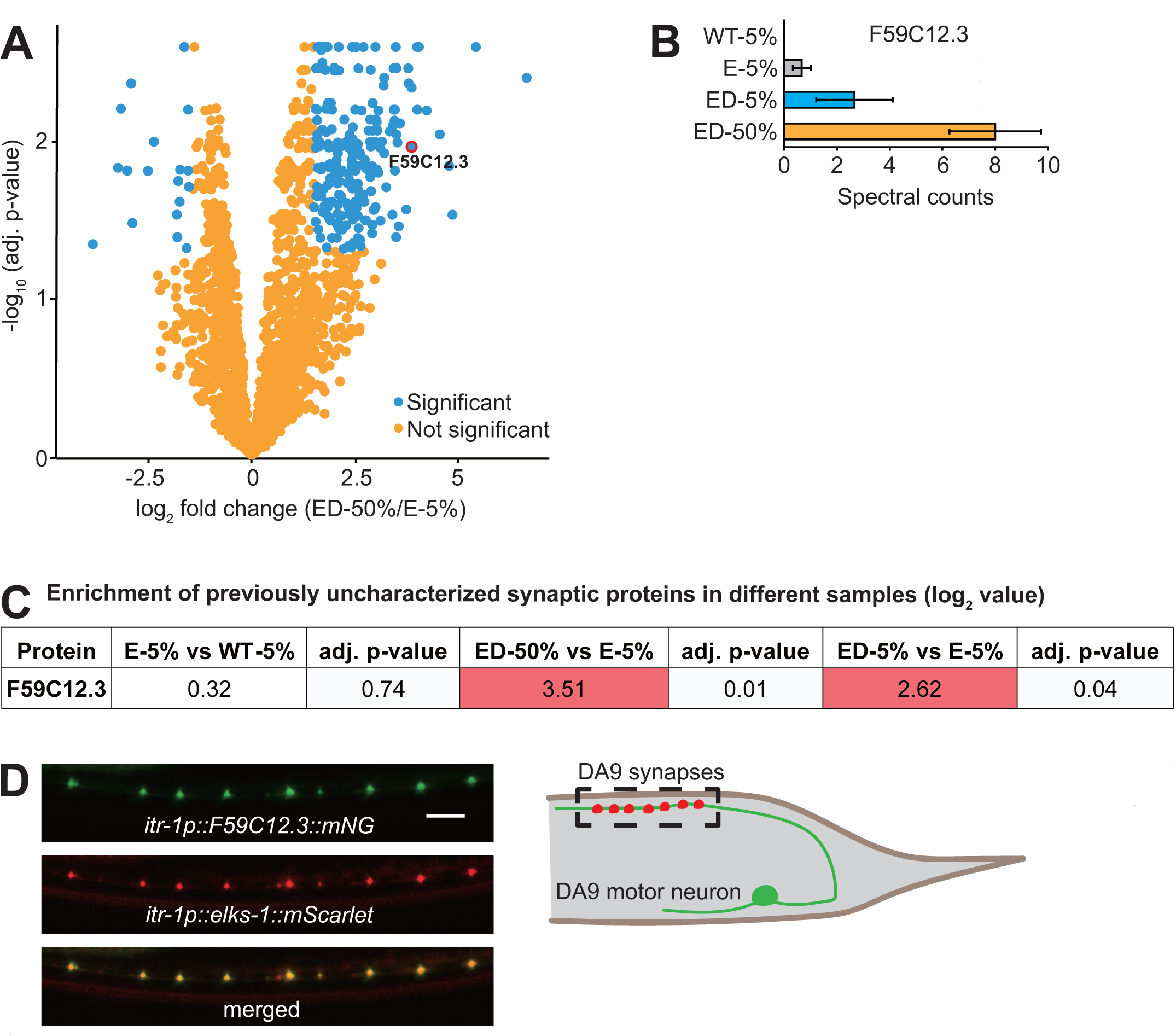
Previously uncharacterized proteins enriched in depleted samples. (**A**) Significantly enriched proteins in the ELKS-1 depleted samples (ED-50%) compared to undepleted ones (E-5%). (**B** and **C**) Mean spectral counts (**B**) and log_2_ fold enrichment (**C**) of previously uncharacterized protein F59C12.3 obtained via mass spectrometry. (**D**) Confocal microscopy images of transgenic *C. elegans* expressing an *F59C12.3::mNG* translational fusion in the DA9 motor neuron. *elks-1::mSc,* used as a control to mark DA9 synapses, confirms these proteins are synaptic. Scale bar: 5 µm.

## Discussion

In *C. elegans* and other species, four endogenously biotinylated carboxylases, MCCC-1/MCCC, PCCA-1/PCCA, POD-2/ACACA, PYC-1/PC (MP3), dominate the biotinylated proteome. Peptides from these large and abundant carboxylases obscure peptides from less abundant biotinylated proteins when samples affinity-purified on streptavidin beads are analyzed by mass spectrometry. Using CRISPR/Cas9 genome editing, we tagged the MP3 proteins with a His_10_ tag, allowing us to deplete them using affinity purification, substantially increasing the sensitivity of MS analysis. We expect depletion will be especially useful when the POI tagged with TurboID is expressed at low levels, or only in a subset of cells. Depleting the carboxylases not only reduces noise by removing confounding peptides, it also increases signal as it allows more of the sought after peptides to be loaded onto the mass spectrometer without saturating the run.

Adding a depletion step to TurboID experiments using the active zone protein ELKS- 1, highlights many proteins known to be synaptically-enriched that are not highlighted when we omit depletion. These proteins include SEL-5/AAK1, an ortholog of human AP2 associated kinase 1 (24); UNC-41/Stonin, which functions with the AP2 complex to recycle synaptic vesicle membranes (25); RIC-8, which localizes to cholinergic synapses along the ventral nerve cord (26); the Aryl hydrocarbon receptor interacting protein (AIP) related protein AIPR-1, which physically interacts with ryanodine receptor at worm synapses (27); UNC-31/CAPS; PAK-1/PAK1; SAD-1/Sad kinase; synaptobrevin SNB-1; SYD-2/Liprin-alpha; NAB- 1/PPP1R9A; GIT-1/GIT1; RIC-4/SNAP23 and TOM-1/STXBP5. MP3 depletion also highlights ELKS-1 interactors whose enrichment at synapses is less well- documented. We confirmed the synaptic localization of one such protein, F59C12.3. F59C12.3 is predicted to be a 548 residue protein orthologous to human AMOT (Angiomotin). The human Angiomotin family is a group of scaffolding proteins associated with tight junctions (28). AMOT-p130, accumulates at postsynaptic densities (PSDs) in rat hippocampal neurons, and interacts with actin, the multiple PDZ domain proteins MUPP1 and PSD-95 (29). Our co-localization data indicate that AMOT also localizes pre-synaptically.

The high binding capacity, simplicity, and low price of IMAC resins are significant advantages to their use for MP3 depletion. There are also some limitations that should be considered before employing this approach. IMAC resins are often incompatible with buffers containing SDS, EDTA, or DTT, all of which were present in the buffers of our original TurboID protocol (This study; 30). SDS is an anionic detergent used to solubilize and denature proteins, and is particularly effective for solubilizing membrane proteins e.g. synaptic proteins. We tested several detergents including CHAPS, Tween-20 and Triton-X100 to avoid using SDS. Although capture of MP3 was effective in the absence of SDS, protein extraction was not nearly as effective as the original SDS buffer (Fig. S2B). To overcome this obstacle, we started the protein purification protocol with 1% SDS and then reduced this high concentration using desalting columns and a dilution step in Ni-NTA binding buffer.

Schlager et al. (30) removed SDS from protein solutions through precipitation by cooling combined with low concentrations of sarkosyl (31). We tested this protocol, but did not achieve an effective level of MP3 depletion (data not shown). In addition to SDS, the reducing agent DTT and chelating agent EDTA also substantially decrease binding of His-tagged proteins to divalent metal-based (mostly Nickel or Cobalt) resins. DTT reduces and EDTA sequesters the metal ions of an IMAC resin, making them unsuitable for IMAC applications. We therefore excluded EDTA and DTT from our buffers. A short peptide-based affinity purification tag, such as FLAG, might be an alternative to His tag to achieve better MP3 depletion and may not require the removal of SDS, EDTA or DTT from extraction buffers. As highlighted previously (12), urea-containing buffers should be prepared fresh – MP3 depletion was significantly impaired when we used old 8M urea Ni-NTA buffer (data not shown). Including low levels of imidazole is usually advised to minimize non-specific binding to IMAC-based columns or resins (32). Although His-tagged protein purity was not our primary concern, we compared binding of His-tagged proteins and other proteins to Ni-NTA resin in the presence or absence of 5 mM imidazole in Ni-NTA binding buffer. We did not observe a noticeable change in the bound proteins as judged by western blot analysis and Coomassie staining, and therefore omitted imidazole from our buffers (data not shown).

In summary, His_10_ tags can be added to endogenously-biotinylated carboxylases without disrupting their essential function. The tagged enzymes can then be depleted efficiently using IMAC, providing a reliable way to improve detection and enrichment of biotinylated proteins in proximity labeling experiments. Here, we used depletion with TurboID labeling, but the method is applicable to proximity labelling experiments that use other enzymes to generate reactive biotin, e.g. APEX2 or HRP. We have successfully employed this protocol to different POIs, ranging from transcription factors to structural proteins, localized in different subcellular compartments e.g. endoplasmic reticulum, cytoplasm, nucleus (data not shown). Carboxylase depletion will improve the reach of proximity labelling to single cells and subcellular structures, and is likely applicable to many model organisms due to the high evolutionary conservation of these carboxylases.

## Materials and methods

### Strains

Worms were grown at room temperature (22°C) on nematode growth medium (NGM) plates seeded with the biotin auxotroph *E. coli* MG1655. *C. elegans* husbandry otherwise followed standard laboratory culture conditions (33). *C. elegans* strains used in this study include:

N2, the wild-type Bristol strain

AX7884 *pod-2(syb1772[pod-2::His_10_]) II; mccc-1(syb1666[mccc-1::His_10_]) IV; pyc- 1(syb1680[pyc-1::His_10_]) V; pcca-1(syb1626[pcca-1::His_10_]) X* quadruple His_10_-tag knock in strain

AX7885 *pod-2(syb1772[pod-2::His_10_]) II; elks-1(syb1710[elks- 1::CeTurboID::mNeongreen::3xFLAG]) mccc-1(syb1666[mccc-1::His_10_]) IV; pyc- 1(syb1680[pyc-1::His_10_]) V; pcca-1(syb1626[pcca-1::His_10_]) X*

AX7897 *dbIs37[rab-3p::mNeongreen::3xFLAG]; pod-2(syb1772[pod-2::His_10_]) II; mccc-1(syb1666[mccc-1::His_10_]) IV; pyc-1(syb1680[pyc-1::His_10_]) V; pcca- 1(syb1626[pcca-1::His_10_]) X*

AX7898 *dbIs24[rab-3p::CeTurboID::mNeongreen::3xFLAG]; pod-2(syb1772[pod- 2::His_10_]) II; mccc-1(syb1666[mccc-1::His_10_]) IV; pyc-1(syb1680[pyc-1::His_10_]) V; pcca-1(syb1626[pcca-1::His_10_]) X*

AX8273 *bpl-1(db1372[bpl-1::degron(AID)]) ieSi57[eft-3p::TIR1::mRuby::unc-54 3’UTR+Cbr unc-119(+)] II*

AX8489 *mccc-1(db1433[mccc-1::degron(AID)::3xFLAG]) IV; pyc-1(db1434[pyc- 1::degron(AID)::3xFLAG]) V; pcca-1(db1437[pcca-1::degron(AID)::3xFLAG]) X; ieSi57[eft-3p::TIR1::mRuby::unc-54 3’UTR+Cbr unc-119(+)] II*

AX8490 *pod-2(db1461[pod-2::degron(AID)::3xFLAG]) II; mccc-1(db1433[mccc- 1::degron(AID)::3xFLAG]) IV; pyc-1(db1434[pyc-1::degron(AID)::3xFLAG]) V; pcca- 1(db1437[pcca-1::degron(AID)::3xFLAG]) X; ieSi57[eft-3p::TIR1::mRuby::unc-54 3’UTR+Cbr unc-119(+)] II*

AX8466 *dbEx1339[itr-1p::F59C12.3::mNeongreen; itr-1p::elks-1 cDNA::mScarlet, myo-2p::RFP]*

### Molecular biology

#### Cloning uncharacterized ELKS-1 interactors

To generate the pPD95.75-mNG and pPD95.75-mSc plasmids the pPD95.75 plasmid was digested with KpnI and BsmI restriction enzymes (NEB), and the linearized plasmid gel-extracted and assembled with PCR-amplified codon-optimized mNeongreen or mScarlet using In-fusion cloning (Takara). To generate pPD95.75-itr- 1p-mNG or pPD95.75-itr-1p-mSc plasmids pPD95.75-mNG plasmid was digested with HindIII and PstI, pPD95.75-mSc with PstI and BamHI restriction enzymes (NEB), the linearized plasmids were gel-extracted and assembled with PCR- amplified *itr-1* promoter using In-fusion cloning.

The ORF for F59C12.3 (∼5.7 kbp) was amplified from *C. elegans* genomic DNA by PCR and cloned into pPD95.75-itr-1p-mNG vector digested with PstI and KpnI by In- fusion cloning. *elks-1* cDNA (∼2.5 kbp) was PCR-amplified from a *C. elegans* cDNA library and cloned into pPD95.75-itr-1p-mSc vector digested with BamHI and KpnI by In-fusion reaction. The resulting expression vectors were injected into the gonad of day 1 adult N2 worms at a concentration of 25 ng/uL.

### CRISPR/Cas9-mediated genome editing

The AX7884 strain was obtained by crossing strains PHX1772 pod-2(syb1772[pod- 2::His_10_]) II, PHX1666 mccc-1(syb1666[mccc-1::His_10_]) IV, PHX1680 pyc-1(syb1680[pyc-1::His_10_]) V and PHX1626 pcca-1(syb1626[pcca-1::His_10_]) X to obtain quadruple His_10_-tag KI strain. PHX1772, PHX1666, PHX1680 and PHX1626 were generated by SunyBiotech upon our request (Fujian, China). AX8273 was generated by engineering bpl-1 so as to add an AID degron to the C terminus of BPL-1. AX8490 was generated by knocking in sequences that encode an AID degron::3xFLAG just before the stop codon of each of the four carboxylase-encoding genes, and using genetic crosses to assemble the four edited genes into a single strain. AX8273 and AX8490 strains were generated according to previously published CRISPR editing protocol (34).

### Primers used for cloning

F59C12.3 genomic DNA (∼5.7 kbp)

**F-F59C12.3-**GGTGGTGGAAGCACACGcATGTATCAGGGAGAGACGAACATTTTAG R- F59C12.3- CTCCCTTCGACACCATGGCAGAAAATTGGTTATCCGCAAGTATTGAC

*elks-1* cDNA (∼2.5 kb)

F-elks-1-cDNA - GGTGGTGGAAGCACACGGGATCCATGGCACCTGGTCCCGCACCATA

R-elks-1-cDNA- CTGCCTCTCCCTTGCTAACCATGGCGGCCCAAATTCCGTCAGCATCG

*itr-1* promoter for pPD95.75-mScarlet plasmid

F-itr-1p-GAAATAAGCTTGCATGCCTGCAGCTATTCCAGAGTTCGTTCCCGAGC R-itr-1p-CCTTTGGCCAATCCCGGGGATCCCGTGTGCTTCCACCACCACTAGC

*itr-1* promoter for pPD95.75-mNeongreen plasmid

F-itr-1p-CAACTTGGAAATGAAATAAGCTTCTATTCCAGAGTTCGTTCCCGAGC R-itr-1p-GATCCTCTAGAGTCGACCTGCAGCGTGTGCTTCCACCACCACTAGC

Codon optimized *mScarlet* for pPD95.75 plasmid

F-mScarlet-GGAGGACCCTTGGAGGGTACCATGGTTAGCAAGGGAGAGGCAG R-mScarlet-CAGTTGGAATTCTACGAATGCTTTACTTGTAAAGCTCATCCATTC

Codon optimized *mNeongreen* for pPD95.75 plasmid

F-mNG-GGAGGACCCTTGGAGGGTACCATGGTGTCGAAGGGAGAAGAGG R-mNG-CAGTTGGAATTCTACGAATGCTCTACTTGTCATCGTCATCCTTG

### Light Microscopy

Confocal microscopy images of transgenic *C. elegans* expressing fluorescent proteins were acquired using a Leica (Wetzlar, Germany) SP8 inverted laser scanning confocal microscope with a 63x 1.2 NA oil-immersion objective, using the LAS X software platform (Leica). The Z-project function in Image J (Rasband, W. S., ImageJ, U. S. National Institutes of Health, Bethesda, Maryland, USA, http://rsbweb.nih.gov/ij/) was used to obtain the figures used in the panels. Animals were mounted on 2% agarose pads and immobilized with 100 µM of sodium azide.

### Auxin-inducible degradation

Auxin-inducible degradation (AID) assays were conducted as described previously (35). Briefly, age-synchronized animals were grown on biotin auxotroph *E. coli strain* MG1655-seeded NGM plates in the presence or absence of 1 mM auxin (IAA, indole-3-acetic acid, Sigma).

### Immunoblotting

Synchronized populations of *C. elegans* grown on *E. coli* MG1655 were harvested at L4 or young adult stage, washed three times in M9 buffer and flash-frozen after adding 4X Bolt^TM^ LDS sample buffer supplemented with fresh DTT. The samples were then thawed, boiled for 10 minutes at 90° C, vortexed mildly for 10 minutes, centrifuged for 30 minutes at 15000 rpm at 4° C and the supernatant collected.

Proteins were transferred to a PVDF membrane (Thermofisher Scientific) following electrophoresis using Bolt 4-12% Bis-Tris Plus gels (Thermofisher Scientific).

Membranes were blocked for 1 hour at room temperature with 1% Casein blocking buffer, and incubated for 1 hour at room temperature with fluorescently-labeled streptavidin, or with HRP-conjugated antibodies. Membranes were then washed 3 times with TBS-T. The following antibodies or protein-HRP conjugates were used for this study: IRDye^®^ 800CW Streptavidin (1:10000 in Casein) (LI-COR Biosciences), Anti-FLAG M2-Peroxidase (1:5000 in 1% Casein buffer) (A8592 Sigma), anti-alpha tubulin-HRP (1:10000 in 1% Casein buffer) (DM1A Abcam ab40742). Membranes were imaged using ChemiDoc the Imaging System (Model MP, Bio-Rad).

### TurboID-based enzymatic protein labeling and extraction of biotinylated proteins from *C. elegans*

Gravid adult *C. elegans* were bleached and the eggs transferred to NGM plates seeded with *E. coli* MG1655 to obtain synchronized populations of worms. The animals were harvested at L4 or young adult stage, washed three times in M9 buffer, incubated at room temperature (22°C) in M9 buffer supplemented with 1 mM biotin, and *E. coli* MG1655 for 2 hours unless stated otherwise. Two hours later the worms were washed three times in M9 buffer and allowed to settle on ice after the last wash. After completely aspirating the M9 buffer, one volume of 2X TBS buffer supplemented with 1 mM PMSF and cOmplete EDTA-free protease inhibitor cocktail (Roche Applied Science) was added to one volume of packed worms then added dropwise to liquid N_2_ to obtain frozen worm ‘popcorn’. A Spex 6875D cryogenic mill was used to grind frozen *C. elegans* to a fine powder which was then stored at - 80°C. Worm powder was thawed in a 50 ml falcon tube while rolling it on a tube roller at room temperature. After the sample was completely thawed, it was centrifuged (1000 rpm, 1 min) to collect the sample at the bottom of the tube. SDS was added to the sample to a final concentration of 1% from a stock solution of 20% SDS. The tubes were gently inverted a few times and immediately incubated at 90°C for 5 minutes. After heat treatment, the samples were sonicated continuously for 1 minute twice, with brief cooling between the two sonication steps. Sonication used a probe sonicator microtip (QSonica 700, microtip 4417, 1.6 mm and an amplitude setting of 50/max). The samples were cooled to room temperature following sonication and adjusted to 2M urea using a stock solution (8M urea, 1% SDS, TBS buffer). The samples were then centrifuged at 60,000 RPM for 30 minutes at 22°C using a benchtop ultracentrifuge Optima MP and MLA-80 rotor (Beckman), and the clear supernatant between the pellet and surface lipid layer transferred to a new tube. Zeba^TM^ spin desalting columns (7K MWCO) (Thermofisher) were equilibrated 3 times with 5 ml TBS buffer containing 2M urea, freshly supplemented with protease inhibitors (Roche cOmplete EDTA-free protease inhibitor cocktail 1 tablet/25 ml; PMSF 1 mM) by centrifugation at 1000g for 5 minutes (or until the buffer was completely eluted from resin). Around 4 ml of clarified sample was then loaded onto the equilibrated spin column and desalted by centrifugation at 1000g for 5 minutes (or until the sample was completely eluted from resin) to remove free biotin. The desalting step was repeated once more using freshly equilibrated columns. Protein concentration in the samples was measured using Pierce^TM^ 660nm protein assay reagent supplemented with Ionic Detergent Compatibility Reagent (IDCR) (ThermoFisher Scientific).

### IMAC-mediated depletion of His-tagged carboxylases

One ml of PureCube Ni-NTA agarose resin slurry (Cube Biotech, Germany) was transferred to a 15 ml Falcon tube, centrifuged for 1 minute at 2000 rpm and the supernatant removed. The Ni-NTA agarose resin slurry was equilibrated twice for 10 minutes in 10 ml of Ni-NTA binding buffer (100 mM NaH_2_PO_4_, 10 mM Tris-HCl, 8M urea, pH 8.0) before mixing it with the protein lysate. One volume of total protein lysate (containing 12.5 mg of proteins) was mixed with three volumes of Ni-NTA binding buffer in a Falcon tube and mixed with equilibrated Ni-NTA resin. The mix was incubated for 2 hours at room temperature using a tube roller. The tube was centrifuged at 2000 rpm for 1 min and the supernatant transferred to a new Falcon tube. Note: extra care must be paid not to transfer Ni-NTA resin along with the supernatant. To confirm efficient carboxylase depletion, the Ni-NTA resin was processed for His-tagged protein purification and western blot analysis.

### His-tagged protein purification

After completely removing protein supernatant in Ni-NTA buffer, the Ni-NTA resin was washed twice by adding 10 ml of Ni-NTA buffer and rotating for 10 minutes on a tube rotator. After the washes, the Falcon tube was centrifuged at 2000 rpm for 1 minute and the buffer aspirated. The resin was transferred to an Eppendorf tube by mixing with 1M imidazole solution (1:1 ratio V:V) for elution of His-tagged proteins from the resin (a small aliquot can be processed to determine carboxylase depletion).

The tube was incubated at room temperature for 30 minutes using a rotator.

The Eppendorf tube was centrifuged at 2000 rpm for 1 minute and the supernatant transferred into a new tube for western blot analysis.

### Streptavidin magnetic bead acetylation

Pierce^TM^ Streptavidin magnetic beads (Thermo Scientific, 100 µl bead slurry per 12,5 mg of *C. elegans* total protein lysate) were washed three times with 1 ml of Buffer 1 (50 mM HEPES-NaOH, pH 7.8, 0.2% Tween-20), briefly incubating them in the buffer and using magnetic separation to retain beads while discarding buffer. Washed beads were resuspended in a mix of 190 µl of Buffer 1 and 10 µl of 100 mM Pierce^TM^ Sulfo-NHS-Acetate (Thermo Scientific, dissolved in DMSO), and incubated for 1 hour at room temperature to acetylate free amines (36). Beads were washed three times with 1 ml of Buffer 2 (50 mM ammonium bicarbonate, 0.2% Tween-20) to stop the reaction.

### Biotinylated protein pulldown and elution

Carboxylase-depleted supernatant was mixed with acetylated magnetic beads in a 15 ml Falcon tube and incubated overnight on a tube roller at room temperature. To collect beads, a neodymium magnet was taped to the side of the Falcon tube and incubated on a rocking platform for one hour, or until all the magnetic beads bind to the magnet. Unbound lysate was aspirated and the beads transferred to a 2 ml LoBind protein tube (Eppendorf) using 2% SDS wash buffer (150 mM NaCl, 1 mM EDTA, 2% SDS, 50 mM Tris-HCl, pH 7.4). Beads were washed twice with 2% SDS wash buffer, once with TBS-T buffer (150 mM NaCl, 50 mM Tris-HCl, 0.2% Tween- 20, pH 7.6), twice with 1M KCl-T wash buffer (1M KCl, 1 mM EDTA, 50 mM Tris-HCl, 0.2% Tween-20, pH 7.4), twice with 0.1 M Na_2_CO_3_-T buffer (0.1 M Na_2_CO_3_, 0.2% Tween-20, pH 11.5), twice with 2M urea-T buffer (2M urea, 10 mM Tris-HCl, 0.2% Tween-20, pH 8.0) and 5 times with TBS buffer (150 mM NaCl, 50 mM Tris-HCl, pH 7.6). During each wash beads were incubated on a rocking platform for 10-15 minutes and transferred to new tubes as often as possible. At the end of this step, beads were kept in TBS buffer and immediately submitted to the mass spec facility for processing.

### Mass spectrometry

Beads were resuspended in 50 µl 1 M urea and 50 mM ammonium bicarbonate. Disulfide bonds were reduced with 2 µl of 250 mM dithiothreitol (DTT) for 30 min at room temperature before adding 2 µl of 500 mM iodoacetamide and incubating for 30 min at room temperature in the dark. Remaining iodoacetamide was quenched with 1 µl of 250 mM DTT for 10 min. Proteins were digested on the beads with 150 ng LysC (Wako Chemicals) in 1.5 µL 50 mM ammonium bicarbonate at 25°C overnight. The supernatant without beads was transferred to a new tube and digested with 150 ng trypsin (Trypsin Gold, Promega) in 1.5 µL 50 mM ammonium bicarbonate at 37°C for 5 hours. The digestion stopped by adding trifluoroacetic acid (TFA) to a final concentration of 0.5 %. The peptides were desalted using C18 Stagetips (37) and separated on an Ultimate 3000 RSLC nano-flow chromatography system (Thermo-Fisher), using a pre-column for sample loading (Acclaim PepMap C18, 2 cm × 0.1 mm, 5 μm, Thermo-Fisher), and a C18 analytical column (Acclaim PepMap C18, 50 cm × 0.75 mm, 2 μm, Thermo-Fisher), by applying a segmented linear gradient from 2% to 35% and finally 80% solvent B (80 % acetonitrile, 0.1 % formic acid; solvent A 0.1 % formic acid) at a flow rate of 230 nL/min over 120 min. Eluting peptides were analyzed on a Q Exactive HF-X Orbitrap mass spectrometer (Thermo Fisher), coupled to the column with a nano-spray ion-source using coated emitter tips (PepSep, MSWil). The mass spectrometer operated in data-dependent acquisition mode (DDA), and survey scans were obtained in a mass range of 375- 1500 m/z with lock mass activated, at a resolution of 120k at 200 m/z and an AGC target value of 3E6. The 20 most intense ions were selected with an isolation width of 1.4 m/z, fragmented in the HCD cell at 28% collision energy and the spectra recorded for a maximum of 120 ms at a target value of 1E5 and a resolution of 30k. Peptides with a charge of +2 to +6 were included for fragmentation, the peptide match and the exclude isotopes features enabled, and selected precursors were dynamically excluded from repeated sampling for 30 seconds.

Raw data were processed using the MaxQuant software package (version 1.6.17.0) (38) and the Uniprot *Caenorhabditis elegans* reference proteome (www.uniprot.org, release 2020_01), as well as a database of most common contaminants. The search was performed with full trypsin specificity and a maximum of two missed cleavages at a protein and peptide spectrum match false discovery rate of 1%. Carbamidomethylation of cysteine residues was set as fixed, and oxidation of methionine and N-terminal acetylation as variable modifications. For label-free quantification the “match between runs” feature and the LFQ function were activated - all other parameters were left at default.

### Data analysis using R scripts

MaxQuant output tables were further processed in R (R Core Team, 2018) (39). Reverse database identifications, contaminant proteins, protein groups identified only by a modified peptide, protein groups with less than three quantitative values in one experimental group, and protein groups with less than 2 razor peptides were removed for further analysis. Missing values were replaced by randomly drawing data points from a normal distribution modeled on the whole dataset (data mean shifted by -1.8 standard deviations, width of distribution of 0.3 standard deviations). Differences between groups were statistically evaluated using the LIMMA package (40) at 5% FDR (Benjamini-Hochberg).

### Proteomics data deposition

The mass spectrometry proteomics data have been deposited to the ProteomeXchange Consortium via the PRIDE partner repository (41) with the dataset identifier XXXXXXX.

### Ethics Statement

The work used the free-living nematode *C. elegans*, for which there is no requirement for review and approval from an institutional animal care and use committee. Transgenic experiments were carried out following ISTA guidelines for such work.

## Acknowledgments

We thank de Bono lab members for helpful comments on the manuscript, and the Mass Spec Facilities at IST Austria and Max Perutz Labs for invaluable discussions and comments on how to optimize mass spec analyses of worm samples. We are grateful to Ekaterina Lashmanova for designing the degron knock-in constructs and preparing the injection mixes for CRISPR/Cas9-mediated genome editing. All LC- MS/MS analyses were performed on instruments of the Vienna BioCenter Core Facilities (VBCF) instrument pool. This work was supported by a Wellcome Investigator Award (209504/Z/17/Z) to MdB and an ISTplus Fellowship to MA (Marie Sklodowska-Curie agreement No 754411).

## Author Contributions

MA and MdB conceived experiments; MA performed experiments; MA and MdB analysed data; MH and WC performed mass spectrometry analysis and processed mass spectrometry data; MA and MdB wrote the manuscript.

---

**Figure S1.**
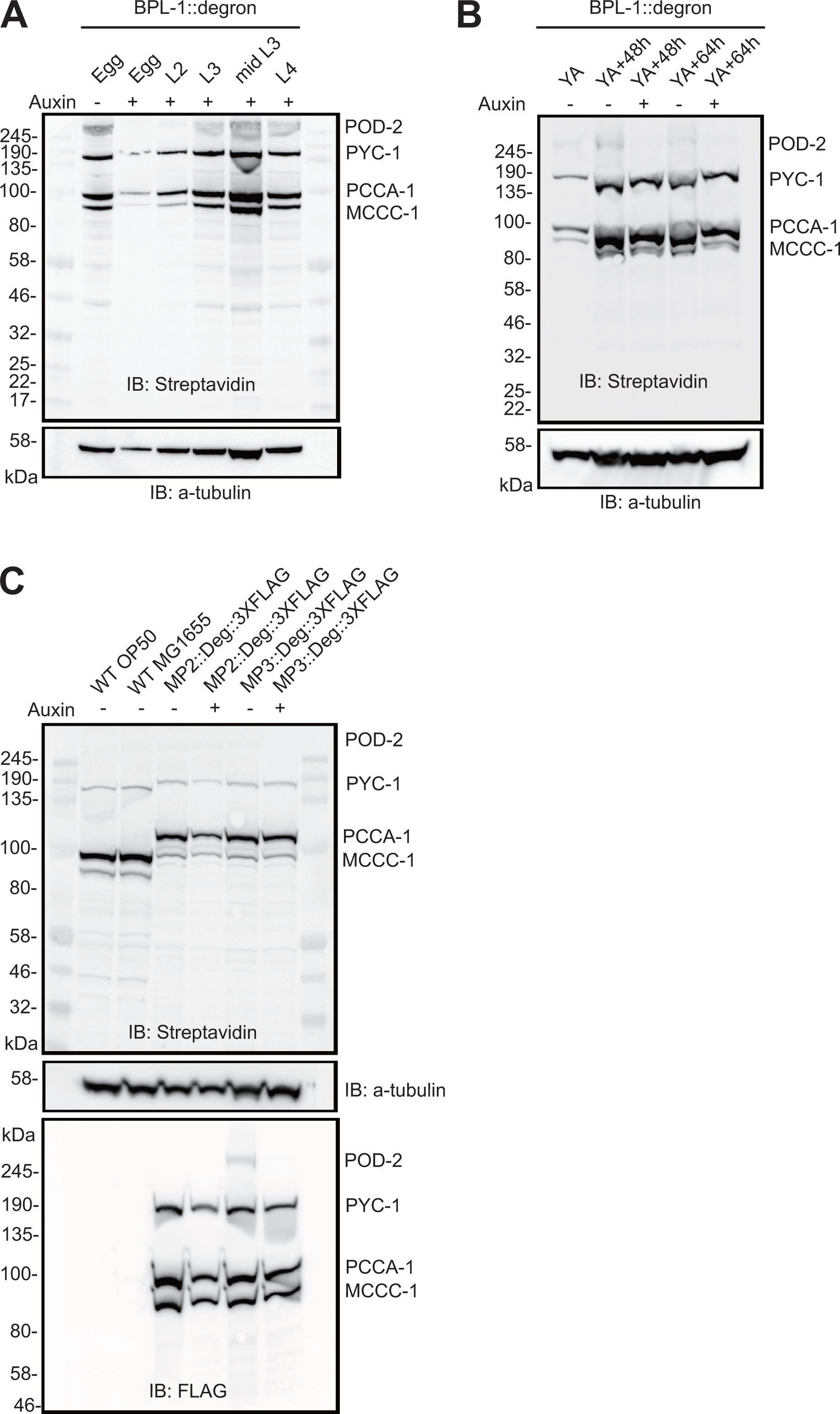
AID-mediated depletion of BPL-1 and MP3. (**A**) The effect of AID-mediated knockdown of BPL-1 starting at different developmental stages on biotinylation of MP3 carboxylases in *C. elegans*. Worms were transferred to auxin plates starting from egg, L2, early or mid-L3 or L4 stages, and harvested as young adults (YA). (**B**) The effect of AID-mediated knockdown of BPL-1 starting at the young adult stage. Worms were transferred to auxin containing plates as young adults and harvested 48 or 64 hours post-YA. (**C**) Depletion of MP2::AID or MP3::AID. A 5-hour incubation on auxin was sufficient to deplete POD-2 but not MCCC-1, PCCA-1 or PYC-1.

**Figure S2.**
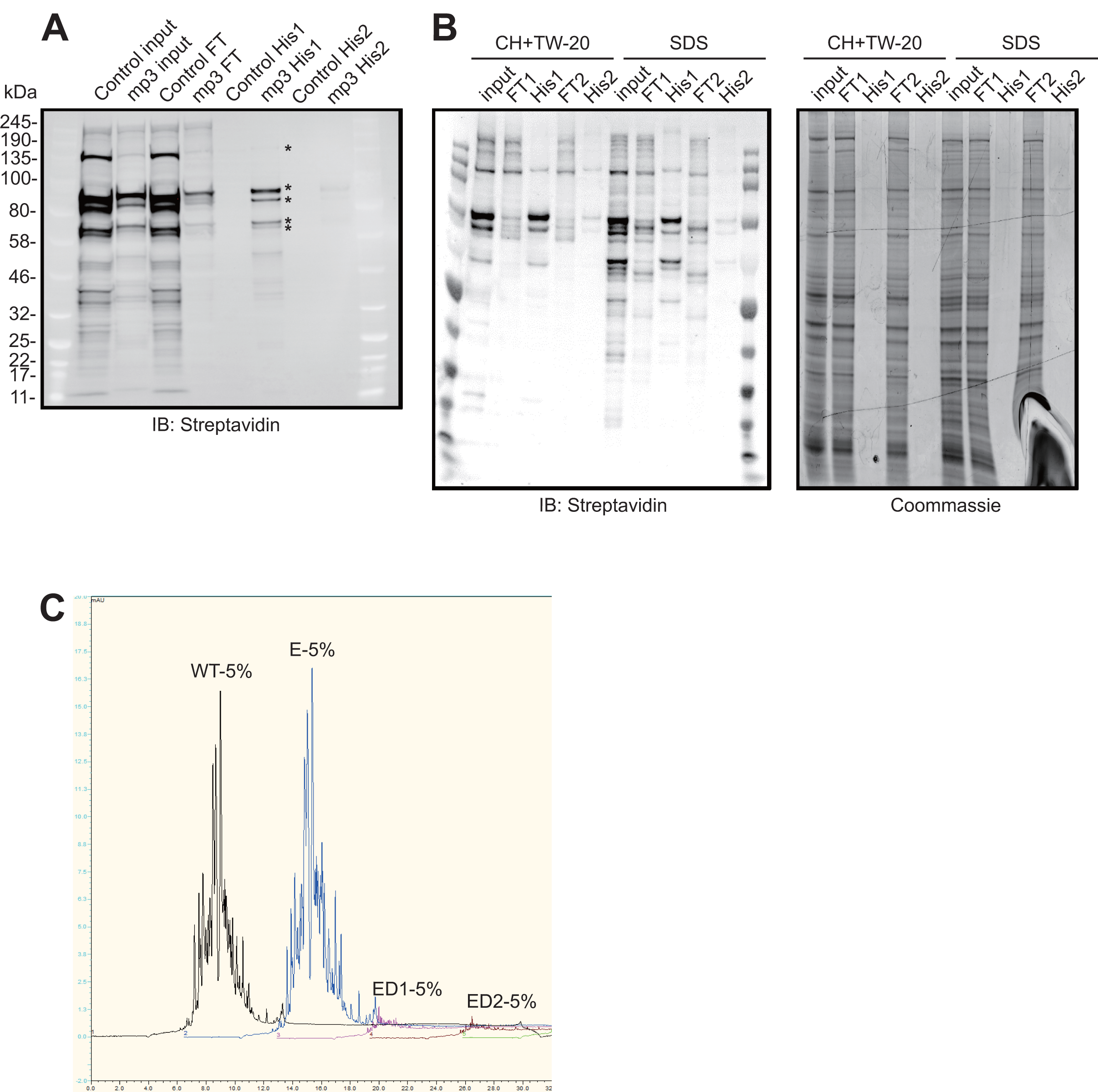
Optimizing carboxylase depletion. (**A**) Ni-NTA resin efficiently binds His_10_- tagged proteins. (**B**) An extraction buffer containing SDS solubilizes the worm proteome more effectively than an extraction buffer containing a mixture of CHAPS, Triton-X100 and Tween-20. (**C**) Ni-NTA depleted and non-depleted worm samples following streptavidin purification and on-bead trypsin digestion analysed using an HPLC-UV system to separate peptides. WT: MP3, undepleted; E: ELKS-1::TbID undepleted; ED1: ELKS-1::TbID samples depleted once; ED2: ELKS-1::TbID samples depleted twice.

**Figure S3.**
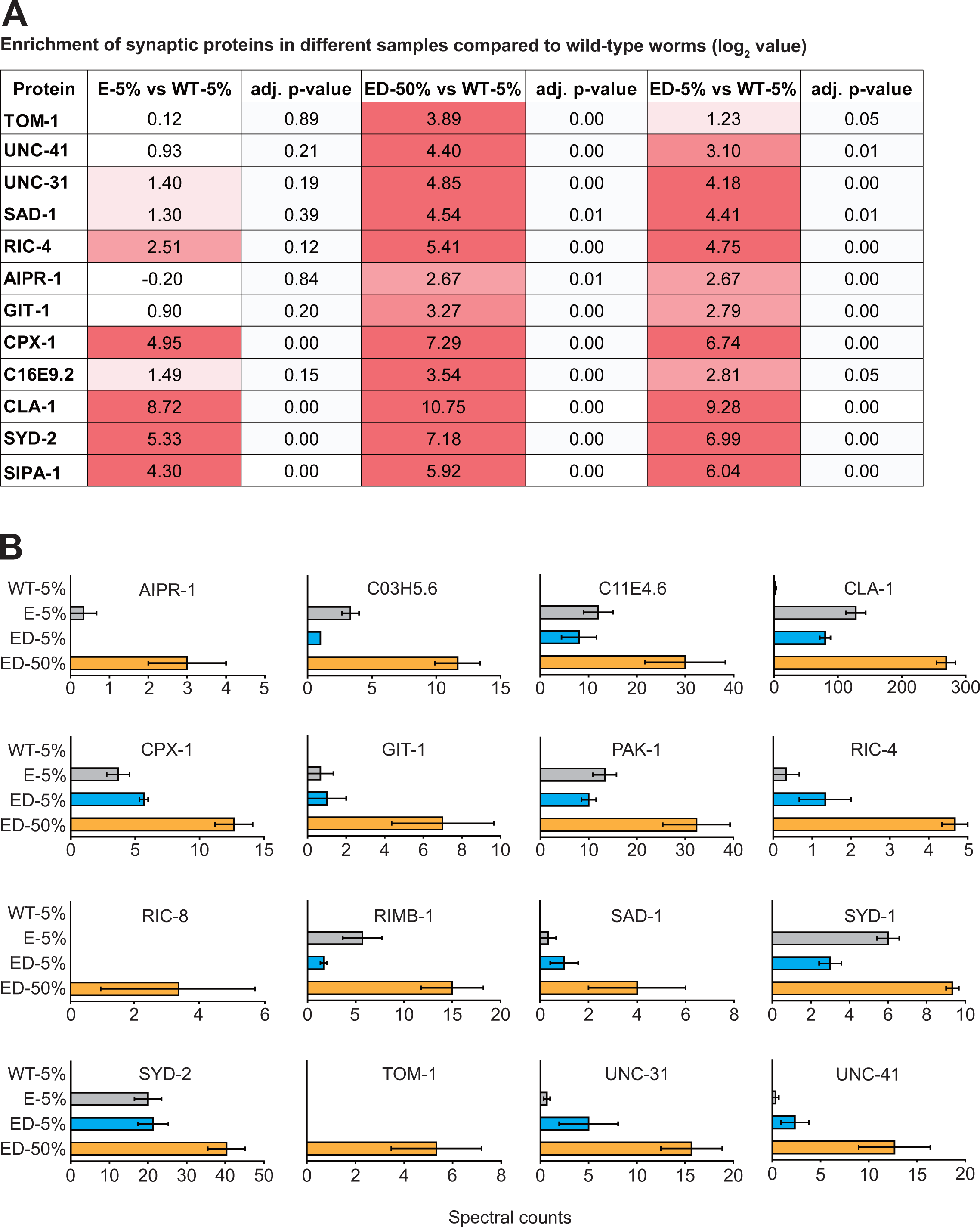
Enrichment of synaptic proteins in depleted and undepleted samples. (**A**) Synaptic proteins in the ELKS-1 undepleted (E-5%) and depleted samples (ED-5% and ED-50%) compared to wild-type samples (WT-5%). (**B**) Mean spectral counts of synaptic proteins in wild-type, undepleted ELKS-1::TbID, and depleted 5% and 50% ELKS-1::TbID samples.

Table S1. LFQ intensity and total spectral counts of all proteins identified in wild-type, undepleted ELKS-1::TbID, and depleted 5% and 50% ELKS-1::TbID samples.

